# Nonstructural protein 1 (nsp1) widespread RNA decay phenotype varies among Coronaviruses

**DOI:** 10.1101/2022.04.19.488803

**Authors:** Yahaira Bermudez, Jacob Miles, Mandy Muller

**Author notes:** equal contribution (Yahaira Bermudez, Jacob Miles).

## Abstract

Extensive remodeling of the host gene expression environment by coronaviruses nsp1 proteins is a well-documented and conserved piece of the coronavirus-host takeover battle. However, whether and how the underlying mechanism of regulation or the transcriptional target landscape differ amongst coronaviruses remains mostly uncharacterized. In this study we use comparative transcriptomics to investigate the diversity of transcriptional targets between four different coronavirus nsp1 proteins (from MERS, SARS1, SARS2 and 229E). In parallel, we performed Affinity Purification followed by Mass-Spectrometry to identify common and divergent interactors between these different nsp1. For all four nsp1 tested, we detected widespread RNA destabilization, confirming that both α- and β-Coronavirus nsp1 broadly affect the host transcriptome. Surprisingly, we observed that even closely related nsp1 showed little similarities in the clustering of genes targeted. Additionally, we show that the RNA targeted by nsp1 from the α-CoV 229E partially overlapped with MERS nsp1 targets. Given MERS nsp1 preferential targeting of nuclear transcripts, these results may indicate that these nsp1 proteins share a similar targeting mechanism. Finally, we show that the interactome of these nsp1 proteins differ widely. Intriguingly, our data indicate that the 229E nsp1, which is the smallest of the nsp1 proteins tested here, interacts with the most host proteins, while MERS nsp1 only engaged with a few host proteins. Collectively, our work highlights that while nsp1 is a rather well-conserved protein with conserved functions across different coronaviruses, its precise effects on the host cell is virus specific.

**Significance:** Coronaviruses extensively co-opt their host gene expression machinery in order to quicky benefit from the host resources. The viral protein nsp1 plays a major role in this takeover as nsp1 is known to induce a widespread shutdown of the host gene expression, both at the RNA and the translational level. Previous work characterized the molecular basis for nsp1-mediated host shutdown. However, this was mostly conducted in the context of β-coronaviruses and in particular SARS-CoV1, CoV2 and MERS due to the important public health burden that these viruses represent. Here instead, we explored the impact of nsp1 on the host using a comparative approach, defining the influence of 4 nsp1 protein from α- and β-coronaviruses. We delineated the impact of these 4 nsp1 on the host transcriptome and mapped their interactome. We revealed that host target range and interactomes vary widely among different nsp1, suggesting a viral-specific targeting. Understanding how these differences shape infection will be important to better inform antiviral drug development.

## Introduction

The past 20 years have seen the emergence of three highly pathogenic human coronaviruses (HCoVs), including severe acute respiratory syndrome (SARS)-CoV, Middle East respiratory syndrome (MERS)-CoV, and SARS-CoV-2. Since 2002 and the first coronavirus epidemic, these viruses have jumped to the forefront of public awareness as major public health threats. Other HCoVs routinely circulate in the human population, such as HCoV-HKU1 or HCoV-229E, which cause mild to moderate upper respiratory tract infections.

Coronaviruses consist of four genera: Alphacoronavirus (α-CoV), Betacoronavirus (β-CoV), Gammacoronavirus (γ-CoV), and Deltacoronavirus (δ-CoV) [1–4]. While γ - and δ-CoVs primarily infect birds [4], α and β-CoV only infect mammals. The three highly pathogenic HCoVs, SARS-CoV (referred to as SARS1 herein), MERS-CoV, and SARS-CoV-2 (referred to as SARS2 herein), belong to the genus β-CoV and are believed to freely circulate in and originate from bats [5,6]. All these viruses have a highly conserved genomic organization and are the largest known RNA viruses to date. The 5’-terminal two-thirds of their genome encodes two overlapping open reading frames (ORF1a and 1b), which results in the production of two large polyproteins. Nsp1 is the most N-terminal peptide released from the ORF1a polyprotein. Nsp1’s role during β-CoV infection has long been studied and nsp1 was revealed to be a host shutoff protein that controls anti-viral responses by globally reducing host gene expression. SARS2 nsp1 similarly to SARS1 nsp1 not only induces translational shutdown [7,8] but also promotes the degradation of its target RNA by binding to the 40S ribosomal subunit [9–11]. Furthermore, MERS-CoV nsp1 selectively targets mRNA synthesized in the host cell nucleus for degradation and thus inhibits translation in host cells [12,13]. How the various nsp1 mediate this extensive shutoff phenotype at the RNA level remains unknown [14,15]. nsp1 shares no resemblance in its primary amino acid sequence or protein structure with any known RNases and has been hypothesized to co-opt a host endonuclease to induce endonucleolytic cleavage of template mRNA transcripts that interact with 40S ribosomes. Yet, the identity of this putative host RNase, while extensively looked for, remains unknown.

Intriguingly, only α and β-CoV encode nsp1, whereas γ- and δ-CoV lack this protein [16,2,17,3,18,19]. The sizes of nsp1 in β-CoV also differ from the α-CoVs nsp1, with the α-CoVs nsp1 being substantially smaller than their β-CoVs counterparts. While these differences may have important consequences on the role of nsp1 during infection, it appears that nsp1 proteins from HCoV-229E and HCoV-NL63 might still be able to bind the 40S ribosomal subunit to affect host mRNA stability[20,21, 17] reminiscing of how the β-CoV nsp1 trigger host shutoff.

The extensive study of the role of SARS2, SARS1 and MERS nsp1 has revealed a pervasive role in reshaping the host gene expression environment. In this study, we set out to compare the impact of nsp1 on the host cell not just from the highly pathogenic β-CoV but also from the α-CoVs 229E. We hypothesized that the extent of nsp1-mediated RNA decay may vary amongst the different HCoV, which could account for some of the severity of these infections. To address this possibility, we generated a library of nsp1-inducible cells from 4 HCoV coronaviruses and explored the extent of nsp1 effect on the host transcriptome by RNA-seq. Interestingly, we found that widespread targeting of RNA is conserved among these nsp1 but the range of targets is different. Moreover, we investigated the interactome of these nsp1 proteins by mass spectrometry to identified common and divergent interactors and identified factors that may contribute to nsp1 targeting of RNA. This work provides important insights into the fundamental differences between highly pathogenic and common coronaviruses nsp1 and refines our understanding of nsp1-meidated decay.

## Results

### Inducible Expression of Coronavirus nsp1 proteins

Previous work has shown conservation in the ability of α- and β-coronaviruses to modulate host gene expression pathways. However, to date, it remains unclear how diverse is the impact on the host transcriptome between α- and β-coronavirus nsp1 proteins. In order to assess the influence of different coronavirus nsp1 proteins on the host gene expression environment, we generated a library of nsp1-inducible cell lines using the nsp1 protein of 4 different HCoV. To generate this library, along with SARS2 and MERS, we constructed 229E and SARS1 coronavirus nsp1 lentiviral plasmids derived from the pLVX-TetOne-Zeo-CoV2-nsp1-3xFlag gifted to us by the Glaunsinger Lab. Following production of lentivirus plasmids, we performed lentiviral transduction using select pLVX CoV nsp1-Flag plasmid and pMD2.G and psPAX2 envelop and packaging plasmids (**Fig. 1A**). After transduction, cells underwent selection using zeocin at a concentration of 325ug/ml. Following selection, to verify that our cell lines were inducible and would produce CoV nsp1 protein, cells were induced with 1ug/ml doxycycline. 24 hours post induction samples were collected and protein samples were western blotted with a Flag antibody (**Fig. 1B**). All four nsp1 proteins express well under induction and at the expected size. Once we verified that all four CoV nsp1 proteins could be properly expressed, we next determined if these induced nsp1 regulated gene expression as expected. As reported before, nsp1 can efficiently degrade GFP transcripts and reduce GFP expression by itself [9,11]. We thus next expressed a GFP reporter, then the transduced cells were either left in an uninduced state, or induced with doxycycline. 24 hours post transfection and induction, GFP expression was measured with fluorescent microscopy (**Fig. 1C**) and quantified using ImageJ (**Fig. 1D**). As expected, induction led to a significant decrease in the intensity of GFP expression in all CoV nsp1 expressing cells. This is in line with the observation that despite being a smaller protein, the 229E nsp1 is able to regulate the expression of genes, likely due to a highly conserved domain [21]. These results show that our library of nsp1 expressing cell lines do not have leaky expression, are inducible, and able to modulate gene expression.

**Figure 1:**
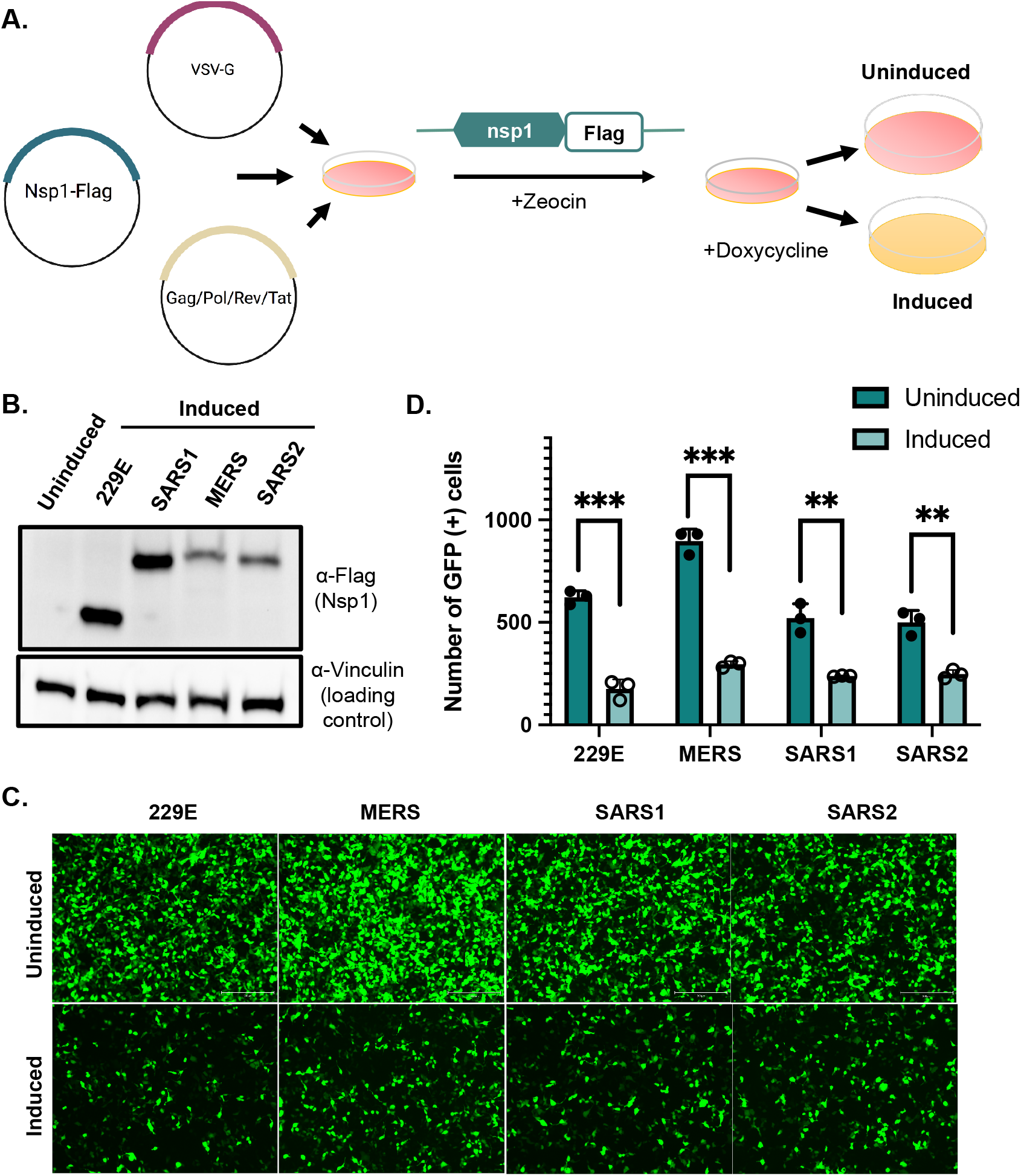
Inducible Expression of 4 Coronavirus nsp1 results in downregulation of GFP. (A) Diagram representing lentiviral transduction and cell induction protocols. HEK293T cells were transfected with pLVX plasmids expressing the 4 nsp1 selected (from 229E, SARS1, SARS2, MERS) under a doxycycline inducible promoter along with the lentiviral envelope and packaging plasmids. (B) Transduced cells were induced with doxycycline for 24h (or left uninduced). Cells were then harvested, lysed, resolved on SDS-PAGE, and immunoblotted with the indicated antibodies. (C) Transduced cells were transfected with a GFP reporter for 24h then nsp1 expression (as indicated) was induced with Doxycycline for 24h. GFP expression was monitored using fluorescent microscopy and quantification of GFP positive cells is shown in panel D. ** P<0.01 *** P<0.001

### Coronavirus nsp1 comparative RNA-seq shows differences in gene expression

After confirming that our inducible cell lines function as expected, we sought to explore the differences in the extent of nsp1 targeting on the host transcriptome. Using the lentivirally transduced cell lines described above, we induced the expression of nsp1 from SARS1, SARS2, MERS, or 229E (using uninduced cells as controls). 24h post induction, total RNA was extracted, polyA selected and cDNA library were prepared and sequenced. Out of a total of 19331 genes identified by RNA-seq, 15779 genes appeared in all 4 datasets (**Supplementary Table 1**). More specifically, we identified 3121 genes that were consistently down-regulated upon expression of all 4 nsp1 and 3833 that were consistently upregulated (**Fig. 2A**). Unsurprisingly, 229E nsp1, as the only representative of an α-CoV, has the most unique pattern of genes up and down regulated, perhaps indicating that its targeting mechanism differs from that of the β-CoV nsp1. Furthermore, fold change patterns induced by 229E and MERS nsp1 as observed by volcano plots were nonstandard as opposed to SARS1 and SARS2 plots (**Fig. 2B**). This might suggest that the fold change distribution may not follow a normal distribution. Previous studies had indicated that MERS preferentially targets nuclear mRNA while our data here represents whole cell RNA pools, which could skew our data representation. This might suggest that 229E nsp1 similarly only targets a subset of RNA in cells. We next performed hierarchical clustering on this comparative transcriptomics dataset. **Figure 2C** shows a heatmap of the correlation matrix across all transcripts. In line with previous analyses, our data indicates that all the nsp1 tested trigger massive RNA degradation with >50% of the detected genes downregulated upon each nsp1 expression (**Fig. 2D**). Based on our analysis, between 30 and 45% of genes had a fold change between 1 and 2, indicating that nsp1 had little to no effect on them. Surprisingly, we also detected close to 15% of genes that seem to be upregulated upon nsp1 induction. Overall, our data indicates that α- and β-coronavirus nsp1 share the ability to widely trigger RNA decay however, the precise target of each of the nsp1 differs, suggesting that the host cell is likely differentially affected.

**Figure 2:**
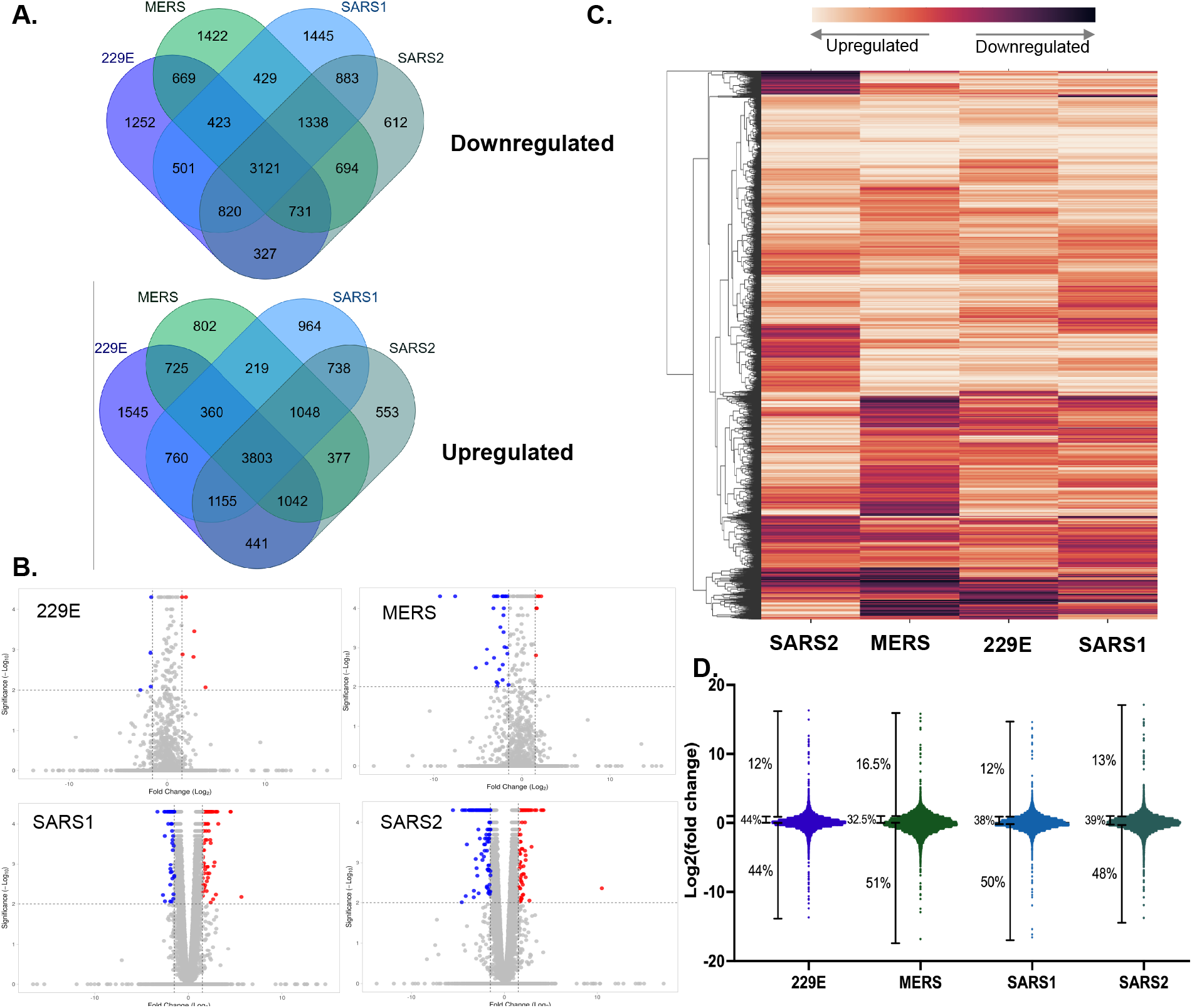
RNA-seq analysis shows differential expression of host genes as impacted by coronavirus nsp1 expression. (A) Venn Diagrams showing the extent of unique and shared genes identified by RNA-seq between each coronavirus nsp1 RNA-seq datasets. (B) Volcano plot of all genes differentially expressed between uninduced samples versus coronavirus nsp1-expressing cells. Data points are shown as a representation of the log2(Fold Change) versus the-log10(p value) plotted by VolcaNoseR2. Significance cut-off was set to the programs default setting, with significantly downregulated genes being labeled blue and significantly upregulated genes being labeled red. (C) Hierachical clustering and heat map of RNA-seq data. Data was clustered based on Fold Change with columns representing expression of different coronavirus nsp1. Expression levels were normalized relative to uninduced sample expression and represented as a heatmap. Transcripts are clustered based on complete linkage method to place genes with high similarities together with dendrogram on the left. (D) Distribution of fold change per nsp1 tested over uninduced sample and corresponding percentages of degraded, unaffected and upregulated transcripts.

### Validation of Coronavirus nsp1 RNA-seq gene expression patterns by RT-qPCR

RNA-seq provided an extensive range of data about the effects of expression of the different coronavirus nsp1s on host gene expression. After data sorting and processing, we noticed some genes with particular patterns of degradation upon expression of the nsp1. We thus next wanted to validate these patterns by RT-qPCR (**Figure 3**). The first gene that we tested was ANKRD1. There are multiples links between ANKRD1 and coronaviruses, for example upon Porcine epidemic diarrhea virus (PEDV) -an α-coronavirus infection-ANKRD1 is downregulated [22], or increased viral load in SARS-CoV-2 infections if ANKRD1 is knock down[23]. Our RNA-seq data showed that ANKRD1 was downregulated upon expression of all tested nsp1. We thus induced the expression of each nsp1 in cells and collected total RNA to examine ANKRD1 expression by RT-qPCR. Our results recapitulate the RNA-seq data. We next tested the gene used chosen INHBE, which showed higher downregulation in SARS2 and MERS, with slight down regulation in SARS1 and 229E in our RNA-seq data. This pattern was again reflected in the RT-qPCR results (**Fig3. B**). We next investigated a gene (SP140L) that appears to be slightly upregulated upon expression of SARS2 and SARS1 nsp1, but downregulated with MERS and 229E nsp1 expression. While our 229E nsp1 in this case appeared to have no effect on SP140L expression, MERS nsp1 expression led to strong downregulation as would be expected from our RNA-seq data (**Fig3. C**).

**Figure 3:**
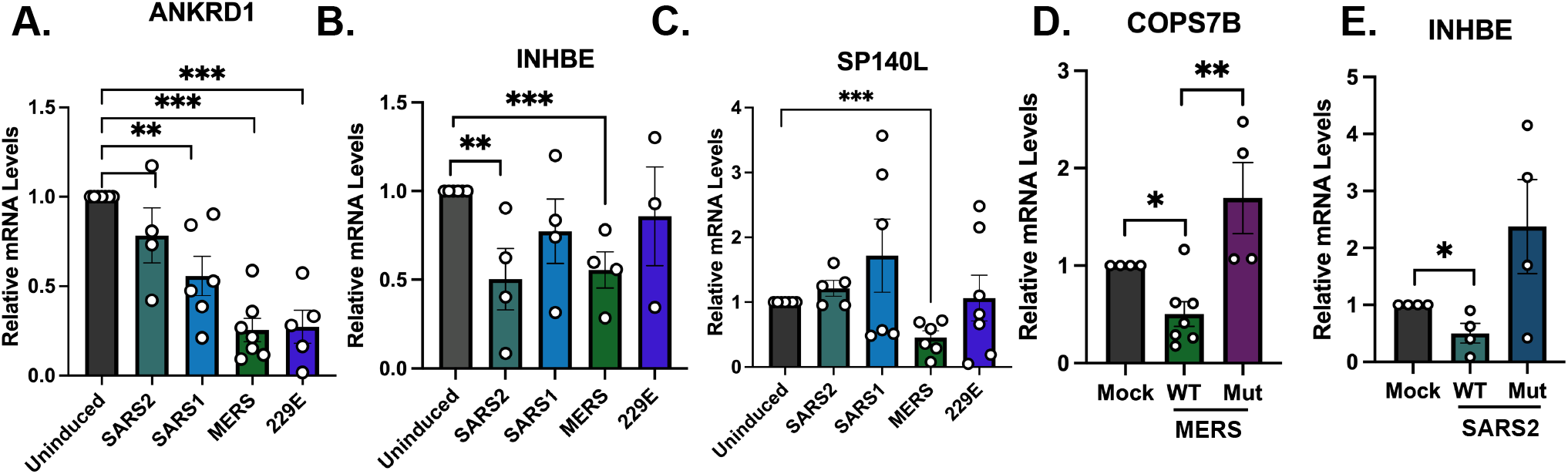
RNA-seq validation. (A-D) 293T cells were either left uninduced or induced for 24hrs as described above, total RNA was harvested and underwent RT-qPCR to measure expression of the endogenous genes ANKRD1 (A), INHBE (B), SP140L (C). (D-E) 293T cells were either left uninduced or induced for 24hrs or transfected with nsp1_mut from MERS (E) or SARS2 (F) as indicated. Total RNA was harvested and underwent RT-qPCR to measure expression of the endogenous genes COP17S or INHBE. n.s., not significant; **, *P*< 0.01; ***, *P*< 0.001.

Our understanding of how nsp1 triggers RNA decay remains mostly vague, however, some nsp1 mutants have been identified to specifically have no RNA decay activity. Residues R146 and K147 in MERS and R124 and K125 in SARS2 have been shown to be required for 40S binding and mutating these residues has been shown to affect RNA turnover [8,24,11,25]. We thus next wanted to validate our RNA-seq data using these mutants. We thus constructed Flag-tagged versions of a MERS nsp1 mutant (R146A+K147A here referred to as MERS nsp1_mut) and a SARS2 nsp1 mutant (R124A+K125A here referred to as SARS2 nsp1_mut), which are both expected to be deficient in RNase activity. For MERS nsp1_mut we looked at COPS7B, which we found to be downregulated by MERS nsp1 expression. Cells were transfected with a WT MERS nsp1, or transfected with the Flag-tagged MERS nsp1_mut. 24h later, total RNA was extracted and used to assess COPS7B expression by RT-qPCR. While WT nsp1 induced a strong reduction in COPS7B expression as expected from the RNA-seq data, the nsp1 mutant failed to induce degradation (**Fig3. D)**. Similarly, for SARS2 nsp1, we compared the expression of INHBE upon WT or mutant expression and saw that nsp1_mut failed to induce INHBE mRNA degradation (**Fig3. E**). Combined together, these results show that all nsp1 tested specifically induce mRNA decay. Additionally, we show that the mutant variants of the SARS2 and MERS nsp1 proteins we have are able to ablate mRNA degradation.

### Nsp1 interactome varies amongst Coronaviruses

Nsp1 contribution to the widespread changes in host mRNA stability is believed to be mediated through protein-protein interaction(s) since nsp1 itself does not appear to act as a nuclease. Yet, very little is known about nsp1 interactions, in particular in the less studied ±-CoV. To better decipher the contribution of nsp1 to the regulation of gene expression, we next compared the interactome of the 4 selected nsp1 using Affinity Purification coupled with LC-MS/MS (AP-MS) to map the nsp1-host protein interaction network (**Fig 4A**). In total, 128 unique proteins were identified in this interactome, of which 74 were unique to 229E nsp1, 8 to MERS nsp1, 21 to SARS1 nsp1 and 9 to SARS2 nsp1 (**Supplementary Table 2**, **Fig 4B**). Notably, among the interactors of nsp1, ribosomal proteins and translation initiation factors were very prominent, which is in line with previous studies investigating nsp1’s interactions as well as consistent with what is known about nsp1 function during infection [26,27]. Furthermore, comparison of the interaction networks highlighted that of the 4 nsp1 tested, nsp1 from MERS was the most divergent (**Fig. 4B**). We next performed a functional analysis of this interactome using Gene Ontology (GO) which revealed 6 functional groups involved in nsp1 mediated-RNA decay and essential cellular biological processes that could promote viral progression (**Fig. 4C, Suppl. table 3**). These results demonstrate the differences between coronavirus nsp1-host proteins interactions and provide insight into how host proteins may contribute to nsp1 role for progression of viral infection.

**Figure 4.**
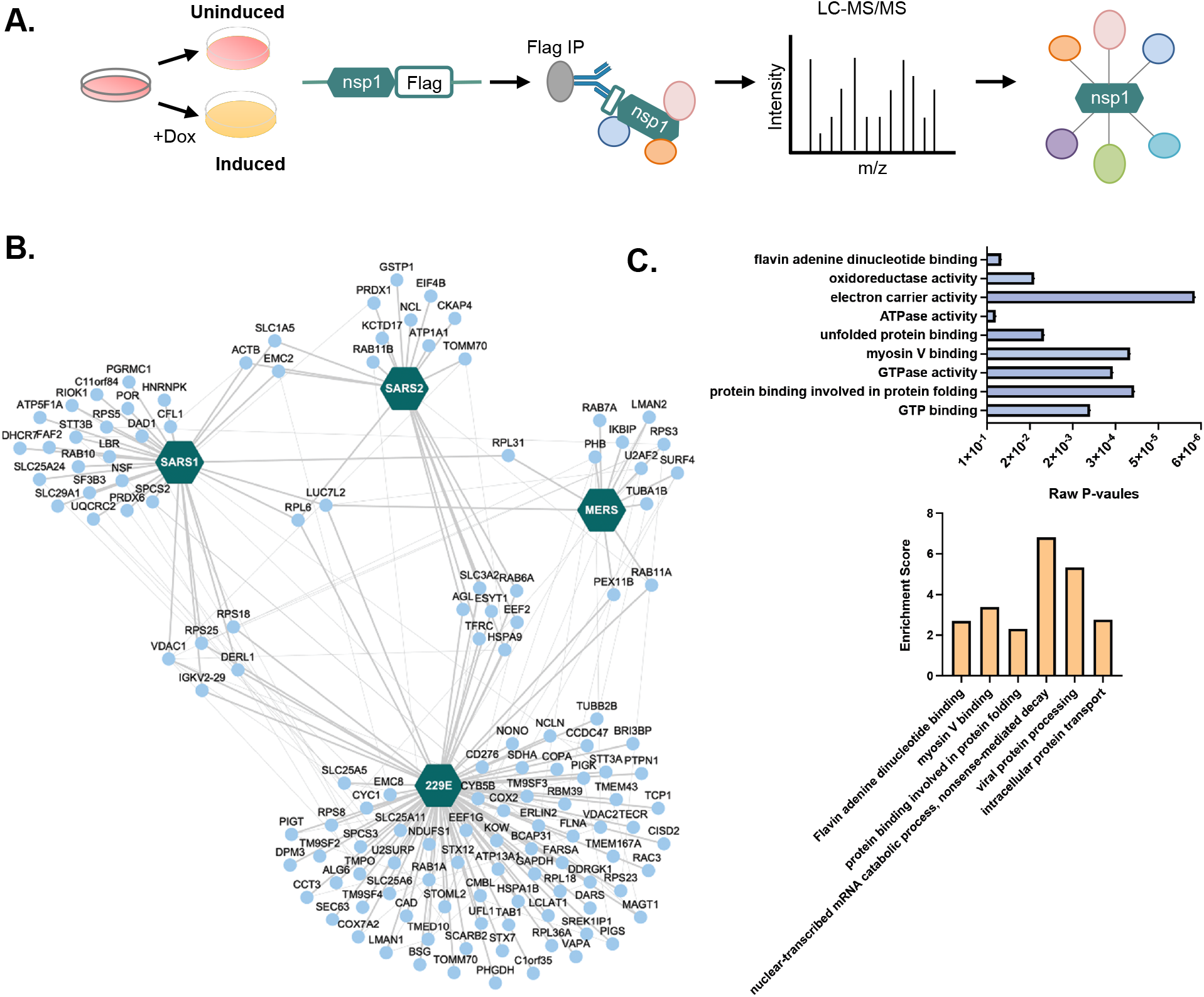
AP-MS of nsp1-host protein interactions shows few shared interactors between coronaviruses. (A) Schematic representation of the Affinity Purification followed by Mass Spectrometry (AP-MS) strategy used to identify the interactors of nsp1 from 4 different coronaviruses. Flag-tagged nsp1 expression was induced in HEK293T as described above. (B) Cytoscape network representation of nsp1-host protein interactions identified by MS. Host proteins (nodes) and the 4 nsp1 (hexagon) are represented. Intra-network interactions amongts host proteins (thin gray lines) were manually curated from the STRING and IntAct databases. (C) Gene Ontology (GO) enrichment analysis was performed on the interacting proteins for each coronavirus nsp1 using DAVID bioinformatic database. The top (blue) histogram shows the raw p-values of most enriched molecular function GO terms and the bottom histogram (orange) represents enrichment scores for the 6 clusters found in the GO term analysis.

## Discussion

Destabilization and degradation of host mRNA is a prevalent strategy employed by numerous viruses as a means of usurping the host gene expression machinery and to dampen the host immune responses[28–36]. Prior research has highlighted the ability of SARS1, SARS2, and MERS coronavirus protein nsp1 to disrupt host gene expression through two prominent methods: modulation of host gene expression through mRNA degradation as well as binding of the 40s ribosomal subunit and subsequent translational arrest [21,22,17]. Moreover, some work also explored the ability of the a-coronaviruses nsp1, such as the one from HCoV-229E, to recapitulate some of the β-coronavirus nsp1 function [21,17]. However, the extent and how conserved nsp1-mediated RNA decay is on the host transcriptome between the different coronavirus nsp1 remains poorly understood. Yet, we know that different nsp1 have different effect on the infected cells. For example, it has recently been shown that SARS2 nsp1 better suppresses STAT1 and STAT2 phosphorylation than the nsp1 protein of SARS1 and MERS thereby more strongly repressing type INF-1 expression[37]. This supports the notion that these proteins may function differently and may show different preferences towards decay and regulation of gene expression. Another example is MERS nsp1 that was shown to preferentially targets transcripts of nuclear origin, while forgoing those transcribed in the cytoplasm[12]. This again highlights how despite having a similar outcome on the host regulation, the mode of action of the various nsp1 may be coronavirus-specific.

Here, we sought to explore the transcriptional landscape of cells expressing the nsp1 of different coronaviruses to better understand the impact of nsp1 between the β-CoVs and the α-CoV 229E. We observed that about half of the transcripts detected by RNA-seq were downregulated upon nsp1 induction, regardless of the origin of nsp1. This suggests that all nsp1 tested in our study have a conserved and likely profound impact on their host cells *via* the regulation of RNA stability. To our surprise, numerous transcripts detected in our sequencing data were non-coding RNAs (ncRNA). Given the prominent association between nsp1 and the ribosome, it has been hypothesized that nsp1 direct role in RNA decay was likely either associated with the host ribosome stalling pathways or alternatively that the endonuclease necessary to mediate nsp1 decay was recruited at the site of nsp1-ribosome interaction. The association of nsp1 and ncRNA would suggest that maybe nsp1 role in RNA decay may be more extensive than previously thought or perhaps that nsp1 can target multiple decay pathways. MERS nsp1 has already been suggested to lead to RNA decay by targeting transcripts independently of ribosome interactions [12,38]. Based on our observation that all the nsp1 tested here target ncRNAs, it could mean that they share this ribosome-independent RNA decay function. Furthermore, it would be interesting to explore this link between nsp1 and host ncRNA, has this could have widespread consequences on the regulation of the host cell.

While SARS1 and SARS2 nsp1 effect on the host transcriptome appear to adopt a “classic” volcano plot shape, indicative of widespread, uniform RNA destabilization on the host transcriptome, MERS and 229E nsp1 expression data sets do not follow this pattern. When we examine the MERS plot, we see many genes undergoing significant negative foldchanges, but also high representation of genes that have significance with little to no foldchange. This could reflect the fact that MERS nsp1 has been suggested to only target nuclear transcribed RNAs, and our RNA-seq data represents total cellular transcripts. As for 229E, however, the vast majority of identified genes represented in the volcano plot showed little significance in foldchange in totality. It could suggest that 229E preferentially target for a subset of transcript, but has only mild effect on the rest of the transcriptome. Alternatively, it has been suggested by Wang *et al*. [17] that nsp1 greatest contribution to host shutoff is not in fact due to RNA decay but more likely because of the translational arrest induced by nsp1 binding to ribosomes. Our data here would indicate that this could be true for 229E nsp1, where we observe less marked RNA decay than its β-CoV counterparts.

Hierarchical clustering of our transcriptomic data highlight that while all nsp1 protein induce large scale RNA decay, there is limited overlap in their specific targets, especially between SARS1 and SARS2 nsp1. Moreover, genes that are heavily downregulated by MERS nsp1 are clustered to the lower half of the heatmap, likely representing nuclear transcripts. This is very similar to the clustered gene degraded by 229E nsp1, suggesting that 229E nsp1 may also preferentially target nuclear transcripts.

We also verified our results using the SARS2 (R124A+K125A) and MERS (R146A+K147A) nsp1 mutants, confirming that these mutants were not able to induce RNA decay like their WT counterparts. As it has been suggested before, while these mutations lead to a loss of mRNA decay potential in the nsp1 protein, this is likely not be linked directly to nsp1 containing RNAse activity. As no nsp1 sequence contains amino acid sequences associated with RNA catalysis, the mutations could potentially disrupt interactions with a protein that does may play this role for nsp1.

Another challenging aspect of studying nsp1 biology is that it has been difficult to identify robust interactions between nsp1 and host proteins. Small and large screens looking either specifically at nsp1 interactome or more globally at the interactome of all CoV proteins have identified only few, rarely confirmed interactors [39,26, 40]. Here, to increase the robustness of our results, we continued with our comparative approach and sought to identify interactors that were shared by these 4 nsp1. Intriguingly, out of this interactome, 229E nsp1 had by far the highest number of unique interactors and MERS nsp1 had the fewest. 229E nsp1, like other α-coronavirus nsp1 are drastically smaller than those found in β-coronaviruses. We can thus speculate that it would require more interacting partners to achieve its primary functions during viral infection. It also possible that 229E nsp1 possess more disordered regions-regions that are known to facilitate protein-protein interactions -as it might be more difficult for a protein of only 111 amino acids to fold into complex structures. Given that MERS nsp1 is known to operate very differently than the nsp1 from SARS1 and SARS2, it is perhaps less surprising to see that its interactome is also very different. While we did not identify in this interactome any host protein with direct RNAse activity that could account for nsp?s role in RNA decay, our gene ontology revealed that many of the interactors that we detected were associated with mRNA catabolic process. It would thus be interesting to explore these interactions and assess whether any of these factors are essential for nsp1 activity.

In this work we sought to explore the similarities and differences between key RNA regulatory proteins during coronavirus infection, to better understand how their expression lead to different viral pathogenic conditions. We observed that even between closely related CoV, there seems to be little similarities in clustering of genes targeted for decay and that there are few protein interactors shared between these different proteins. However, gene targeting for these proteins does not appear to be random, based on Hierarchical clustering the nsp1 seem to target different gene clusters.

---

**Supplementary Table 1. List of genes detected by RNA-seq.**

**Supplementary Table 2. List of mass spectrometry high confidence hits by nsp1.**

**Supplementary Table 3. Go-term analysis on nsp1 interactors.**

## Acknowledgments

We thank all members of the Muller lab for helpful discussions and suggestions. Special thanks to Dr. Britt Glaunsinger and Dr. Ella Hartenian for the pLVX plasmids. We would also like to thank Dr. Ravi Ranjan at the genomics facility for their guidance with sequencing and Dr. Stephen Eyles at the UMASS IALS Mass Spectrometry facility for their help with protocol development and data acquisition.

## Materials AND Methods

### Plasmids and Plasmid Construction

The pLVX-TetOne-Zeo-SARS2-NSP1-3xFlag and pLVX-TetOne-Zeo-MERS-NSP1-3xFlag, pcDNA4 CoV2 nsp1 m1 3xFlag and pcDNA MERS nsp1 m1 3xFlag were a kind gift from the Glaunsinger Lab. CoV-229E sequences were obtained as gBlocks from IDT and subcloned into these plasmids using InFusion cloning (Takara).

### Cells, Transfections, and Lentiviral Transduction

HEK293T cells (ATCC) were grown in Dulbecco’s modified Eagle’s medium (DMEM-Invitrogen) supplemented with 10% fetal bovine serum (FBS). To establish the lentiviral cell lines, HEK293T cells were co-transfected with their respective zeocine-resistant, lentiviral vector along with pMD2.G and psPAX2 the envelop, packaging, and accessory plasmids in DMEM 0% FBS with using Polyjet (SignaGen). 48 hours post transfection the supernatant from transfected cells was collected and filtered through a 0.45 μM filter diluted with serum free media with polybrene at a concentration of 8μg/ml. Mixture was added to fresh HEK293T cells in a 6-well plate and underwent spinfection at 1500rpm for 1.5 hours. Cells were incubated overnight then split the following day into a 10cm plate with media containing 325μg/ml of zeocin. After selection the established lentivirally infected cell lines were maintained in 162.5μg/ml zeocin. For DNA transfection, HEK293T cells were plated and transfected after 24h when 70% confluent using PolyJet (SignaGen).

### Western Blotting

Cell lysates were prepared in TP150 lysis buffer (NaCl, 150mM; Tris, 50mM; NP-40, 0.5%; dithiothreitol [DTT]. 1mM; and protease inhibitor tablets) and quantified by Bradford assay. 20ug of each sample were resolved by SDS-PAGE and Western blotted with following antibodies in TBST (Tris-buffered saline, 0.1% Tween 20): mouse anti-Flag at 1:1000 (Invitrogen) and rabbit anti-Vinculin at 1:2000 (Invitrogen). Primary antibody incubations were followed by horseradish peroxidase (HRP)-conjugated goat anti-mouse or goat anti-rabbit secondary antibodies (1:5000; Southern Biotechnology)

### RT-qPCR

Total RNA was harvested using TRIzol according to the manufacture’s protocol. cDNAs were synthesized from 1 μg of total RNA using AMV reverse transcriptase (Promega) and used directly for quantitative PCR (qPCR) analysis with the SYBR green qPCR kit (Bio-Rad). Signals obtained by qPCR were normalized to those for 18S.

### RNA-seq

6 biological replicates of each lentiviral transduced cells (SARS1, SARS2, MERS, 229E and uninduced sample as a control) were grown to 80% confluency then induced with 1ug/ml of Doxycycline (BD Biosciences). After 24hr cells were collected in TRIzol Reagent and RNA was harvested following the manufacturer’s protocol. Purity of samples were analyzed via bioanalyzer. Following poly(A) selection, libraries underwent 76-base paired-end sequencing using the NextSeq500 Mid-150 cycle kit on a NextSeq 500. Read quality was assessed using fastqc. Using Galaxy[41] reads were then aligned to the human genome (hg38) by Bowtie2 and differential expression analysis were performed using Cufflink and Cuffdiff[42]. For graphical representation in the heatmap, fold change values were saturated by a hyperbolic tan function with a cutoff set at 10. Hierarchical clustering was generated in Python using the SciPy package with complete linkage and Euclidian distance. Volcano plots were developed using the program VolcaNoseR (LCAM) by plotting the log_2_Fold Change vs-log_10_p-value, with significance thresholds left at the default settings and the top 10 most significant genes highlighted and labelled. Comparison of overlapping genes and generation of Venn diagrams were generated using Multiple List Comparator (Molbiotools).

### Immunoprecipitation

Cells were lysed in a low-salt lysis buffer (150 mM NaCl, 0.5% NP-40, 50 mM Tris [pH 8], 1 mM DTT, and protease inhibitor cocktail), and protein concentrations were determined by Bradford assay. For FLAG construct pull-downs, 400 μg of total protein lysates were incubated overnight with Anti-FLAG M2 Magnetic Beads (Sigma) or control G-coupled magnetic beads. Beads were then washed extensively with lysis buffer. Lastly, samples were resuspended in 4X laemmli loading dye before resolution by SDS-PAGE and further Western blotting.

### Mass Spectrometry

Lentiviral transduced cells were grown in 10 cm plates to an 80% confluency and then induced with 1μg/ml of Doxycycline (BD Biosciences). 24 hours post-induction, cells were harvested and lysed, and immunoprecipitation (as mentioned above) was performed overnight at 4C. Samples were extensively washed, and trypsin digested overnight. Samples were then cleaned up using a C_18_ column and mass spectral data obtained from the University of Massachusetts Mass Spectrometry Center using an Orbitrap Fusion mass spectrometer. Raw data was filtered based on the number of peptides for each hit and Gene Ontology (GO) enrichment analysis was performed on the human interacting proteins for each coronavirus using DAVID bioinformatic database. Top enriched and shared clusters are identified on the network using the Cytoscape software

### Statistical analysis

All results are expressed as means ± standard errors of the means (SEMs) of experiments independently repeated at least three times (individual replicate points are shown on bar graphs). Unpaired Student’s *t*est was used to evaluate the statistical difference between samples. Significance was evaluated with *P* values as indicated in figure legends.

